# Interlimb coordination in Parkinson’s Disease is minimally affected by a visuospatial dual task

**DOI:** 10.1101/2022.07.15.500215

**Authors:** Allen Hill, Julie Nantel

**Affiliations:** School of Human Kinetics, University of Ottawa, Ottawa, ON, Canada

**Keywords:** Parkinson’s disease, dual task, interlimb coordination, relative phase, more affected side

## Abstract

Parkinson’s disease (PD) leads to reduced spatial and temporal interlimb coordination during gait as well as reduced coordination in the upper or lower limbs. Multi-tasking when walking is common during real-world activities, and affects some gait characteristics, like gait speed and variability. However, the impact of a dual task (DT) on intra and interlimb coordination of both lower and upper limbs when walking in people with PD remains unknown. Seventeen volunteers with mild to moderate PD (11 males, 65 ± 8 years, 173 ± 8 cm, 74 ± 20 kg, Unified Parkinson’s Disease Rating Scale motor section 10 ± 5) participated in gait trials in an Extended-CAREN system, which includes a treadmill, 12-camera Vicon motion capture system, and a 180° field-of-view virtual reality projection screen. Participants completed a 3 min walking trial and a 2 min visuospatial word recognition DT trial at their preferred walking pace. Single and DT were compared with a paired t-test, and the less and more affected (LA, MA) sides were tested for equivalence in sensitivity to the DT. During the DT, we found the LA shoulder ROM decreased by 1.5°, and the LA shoulder peak flexion decreased by 1.1° (p<.028, g_av_>.12). The LA and MA hip ROM were differently affected by the dual task (p=.023), and intralimb coordination was affected by dual tasking equivalently between sides (p=.004). These results suggest that during normal single-task gait, people with PD use attentional resources to compensate for reduced arm swing. Furthermore, our results indicate that any effect of DT on lower intralimb coordination is not meaningfully different between the LA and MA sides.

**Statements and Declarations:** The authors have no relevant financial or non-financial interests to disclose.

## Introduction

Parkinson’s disease (PD) is a multisystem neurodegenerative disease characterized by the loss of dopaminergic neurons in the substantia nigra with cascading effects in other regions, including those involved in cholinergic systems (Yarnall et al., 2011; Poewe et al., 2017). The neurodegeneration in PD begins unilaterally and progresses bilaterally, and motor symptoms mirror this unilateral emergence and bilateral progression, commonly resulting in one side being more affected than the other (Djaldetti et al., 2006). In addition to the three cardinal motor symptoms of PD—bradykinesia, rigidity, and tremor— people with PD develop further motor deficits, such as hypokinesia (reduced movement amplitude), increased movement variability, gait asymmetry, and postural instability, all of which may contribute to or reflect impairments in coordinating the upper and lower limbs during locomotion (van Emmerik & Wagenaar, 1996; Yogev et al., 2007; Plotnik et al., 2007; Mirelman et al., 2019). Coordination of the upper and lower limbs is required to maintain dynamic gait stability in the presence of major and minor perturbations (Marigold & Misiaszek, 2009; Krasovsky et al., 2012). In addition, gait tasks involving speed modulation require adaptations in interlimb coordination. Such tasks have shown kinematic differences in older adult fallers compared to non-faller peers (Barak et al., 2006; Shishov et al., 2017).

Coordination is defined as the context- and phase-dependent control of spatial and temporal cyclical relationships between body segments (Krasovsky & Levin, 2010). One prominent feature of normal gait coordination is the frequency-matched anti-phase swing (180 deg phase offset) between the arms, legs, and ipsilateral arm-leg pairs, while contralateral arm-leg pairs swing in phase with each other (Wagenaar & van Emmerik, 2000). (One-to-one arm and leg swing frequencies are expected for gait speeds above 0.8 m/s, while slower gait sometimes exhibits 2:1 arm swing to stride frequencies (Wagenaar & van Emmerik, 2000).) These phase relationships result in gait which is generally temporally and spatially symmetric in healthy adults (Sadeghi et al., 2000; Killeen et al., 2018). In contrast, asymmetry in spatial or temporal gait characteristics is recognized as a disruption in the coordination of normal gait and occurs in several pathological populations, including those with PD (Yogev et al., 2007; Huang et al., 2012; Park et al., 2016).

One approach to measuring coordination is based on dynamical systems theory and uses the continuous relative phase (CRP) between coupled oscillators to characterize the state of the system (Haken et al., 1985). Previous studies using CRP have found that, compared to healthy peers, people with PD have reduced coordination stability and increased average phase error (from anti- or in-phase) for upper and lower interlimb coordination during gait and for upper limb coordination during bimanual tasks (van Emmerik & Wagenaar, 1996; Winogrodzka et al., 2005; Almeida & Brown, 2013). Other studies support the presence of reduced interlimb coordination stability in PD (Roemmich et al., 2013), and increased phase error in some upper and lower limb pairs compared to healthy peers and comparing PD freezers to non-freezers (Nanhoe-Mahabier et al., 2011).

Homologous limb coordination is decreased in PD as well. Plotnik and collaborators (2007) reported worse lower limb coordination in people with PD compared to healthy controls using the phase coordination index (PCI), a combined measure of accuracy and consistency in the anti-phase coordination of step timing during gait. Coordination between the upper limbs is also reduced or altered in people with PD compared to healthy controls (Huang et al., 2012; Sterling et al., 2015).

Although there is support for coordination deficits in PD, the more and less affected (MA and LA) sides may contribute differently to interlimb coordination amidst the predominantly unilateral symptoms of early PD. However, the relative nature of coordination makes it difficult to separately characterize coordination between the MA and LA sides. Some studies have found that the LA side maintains an ability to adapt to and compensate for task requirements, such as split-belt walking (Roemmich et al., 2014) or dual-tasking during gait (Siragy & Nantel, 2020); despite the potential benefits of adaptation, this may appear as “worse” coordination due to the decreased similarity between sides. Other studies have found that people with PD showed increased PCI (ie less accurate and/or consistent coordination) when asked to use non-preferred step lengths and times, or when asked to adapt to external mechanical constraints, such as walking on split-belt treadmills (Williams et al., 2013; Fasano et al., 2016); the authors suggest that the increased PCI indicates a generally reduced ability to adapt (lower limb) coordination in PD. However, adaption to split-belt walking differed between MA and LA sides, depending on which side was sped up (Fasano et al., 2016), and therapy focused on the LA side may be more effective than a balanced or MA-side focus (Ricciardi et al., 2015). In addition, two studies have found that ipsilateral arm-leg coordination is reduced on the MA side (Nanhoe-Mahabier et al., 2011; Roemmich et al., 2013). CRP can be used to measure coordination stability within individual limbs (i.e. intralimb coordination), which could contribute to a better understanding of coordination deficits in specific limbs (Barela et al., 2000; Byrne et al., 2002). However, we are not aware of previous studies having investigated this in people with PD.

Performing a secondary task during gait adds an attentional stressor that can reveal the loss of automaticity for some aspects of gait (e.g. gait speed, gait variability), and this is measured as a change in performance while dual tasking, often referred to as the dual-task cost (Yogev-Seligmann et al. 2008). In addition, experimental studies have shown that coordination of bimanual movement patterns can be stabilized by an attentional focus, but a dual task serving as an attentional distractor interferes with the stabilization (Temprado et al., 1999). Since gait while dual tasking may be more similar to gait in the real world than typical laboratory recordings of steady state gait (Hillel et al., 2019), dual tasking during gait may offer an ecologically valid method of manipulating attention to highlight coordinative deficits. Studies of coordination and dual tasking in people with PD show a larger decline in lower limb coordination, measured by PCI, during a dual task compared to healthy controls, and PCI in PD fallers and freezers is more affected by a dual task than PD non-fallers and non-freezers (Plotnik & Hausdorff, 2008; Plotnik et al., 2009; Plotnik, Giladi, et al., 2011). In addition, studies have found reduced arm swing during gait in people with PD during a dual task compared to single task gait (Mirelman et al., 2016; Baron et al., 2018). These results suggest that loss of automaticity in coordination may be associated with the progression of PD to more severe outcomes.

While effects of dual tasking on bilateral coordination and asymmetry have been found (Plotnik, Dagan, et al., 2011), as well as differences in spatiotemporal and movement characteristics between the MA and LA limbs in the upper and lower body (Mirelman et al., 2016; Siragy & Nantel, 2020), the effect of motor symptom presentation (i.e. MA and LA sides) in coordination within and between the upper and lower limbs has not been sufficiently explored. Differential dual-task cost between sides in interlimb coordination in people with PD could provide evidence of how the asymmetric presentation of the disease affects coordination in the upper and lower limbs. Therefore, the purpose of this study is to investigate MA/LA side differences in coordination deficits in people with PD by observing interlimb coordination within and between the upper and lower limbs during single and dual-task gait.

## Methods

### Participants

Volunteers with mild to moderate PD (between I-III Hoehn & Yahr) were recruited from the Ottawa-Gatineau area. Exclusion criteria included any additional neurological impairment, a recent orthopedic injury or surgery that could interfere with gait, the use of a walking aid, or any discomfort with using a projected virtual reality system. Participant characteristics for sex, age, height, weight, handedness, Unified Parkinson’s Disease Rating Scale (UPDRS) motor section (III), freezing and falling status, and interval since diagnosis of PD were recorded. The MA side was defined as the side where PD motor symptoms first occurred, as reported by participants. Falling status was self-reported based on conservative criteria of having fallen at least once in the past year, including non-injurious falls. Participants were tested when optimally medicated (“ON”) by their normal medication. All participants provided written informed consent, and the study was approved by local ethics review boards.

### Protocol

Data was collected using the CAREN system (CAREN-Extended, Motekforce Link, Amsterdam, NL). The CAREN system consists of an instrumented split-belt treadmill (Bertec Corp., Columbus. OH) embedded in a six degree-of-freedom motion platform, a 12-camera motion capture system (Vicon, Oxford, UK), and a 180 deg field of view projection screen. The participants wore a safety harness attached to an overhead frame on the motion platform to prevent falls without restricting movement.

Participants were allowed an initial familiarization period with the CAREN system for several minutes until they reported being comfortable; the familiarization period was also used to determine their preferred walking speed for the gait trials. The single task trial was 3 min long, while the dual task trial was 2 min long. The dual task consisted of a visuospatial word recognition and acknowledgement task where a word was shown at eye level at a random position between 20-70 deg to the left or right of center; participants were asked to notice and acknowledge the word by reading it aloud. Twelve words were randomly drawn from a standard list of 16 possible words in the native language of the participant (English or French). The dual task began 20 s into the trial, and a new word was shown for 3 s every 2-4 s for 80 s. The dual task was designed to be an ecologically valid recreation of common daily life situations (e.g. public transportation terminal, etc.) requiring perception and comprehension of visual cues (Siragy & Nantel, 2020; Ahmadi et al., 2021). The dual task trial occurred after the single task trial. Participants were allowed to rest between trials when requested.

### Data reduction

A set of 57 markers (Wilken et al., 2012) was used to capture full-body kinematics at 100 Hz; marker data was then filtered using a 4th order dual-pass Butterworth low-pass filter with a cutoff frequency of 12 Hz. Low pass filter cutoff frequency was chosen based on a residual frequency analysis of marker data using an RMS noise of 0.5 mm measured from static markers fixed to the motion platform (Winter, 2009). Gait events were calculated using an algorithm based on the local extrema of the vertical position and velocity of the heel marker (Roerdink et al., 2008). OpenSim was used with the Rajagopal et al. (2016) model to perform inverse kinematics to extract bilateral knee, elbow, shoulder, and hip flexion/extension angles (Delp et al., 2007). The wrist pronation/supination range of motion of the model was expanded from 90 deg to 160 deg to better match normative biomechanical characteristics (Shaaban et al., 2008).

To detect changes in spatial coordination, range of motion (ROM), ROM coefficient of variation (COV), and peak flexion were measured on the MA and LA sides for the shoulder and hip joints. Similarly, changes in temporal coordination were assessed using average intercycle phase variability of CRP (interpreted as coordination stability) to measure for ipsilateral shoulder and hip interlimb joint pairs and intralimb (shoulder and elbow, hip and knee) joint pairs. Continuous phase was calculated as the angle of the complex analytic signal produced by the Hilbert transform after centering the original signal (Lamb & Stöckl, 2014).

### Statistics

Effects of dual tasking on individual variables were tested using paired t-tests. Additionally, a TOST procedure was used to test for equivalence between sides for changes due to dual tasking (DTC); this was done for all lateral variables (e.g. shoulder ROM, which was measured on both LA and MA sides) (Lakens, 2017). Dual task cost was defined as the difference between single and dual task for a given variable. All tests were performed with *α* = 0.05. Unless otherwise noted, equivalence bounds were set per variable using unstandardized values (i.e. the original units/scale, without normalizing to group error) to reduce bias (Lakens, 2017) and estimated from previously published results for healthy older adults. The maximum bilateral difference in DTC for shoulder ROM and peak flexion was estimated to be 3.5° (Plate et al., 2015, tbl. 1; Killeen et al., 2018, fig. 2). The maximum bilateral difference in DTC for shoulder ROM COV was estimated at 6° (Mirelman et al., 2016, tbl. 2). Bounds for hip ROM and peak flexion were set as 0.5°, based on changes in DTC between left and right limbs in PD (Ribeiro et al., 2019, tbl. 2). Lower intralimb phase variability bounds were set to 0.85° (Ghanavati et al., 2014, fig. 2A). Appropriate reference data could not be found for the remaining variables (hip ROM COV, ipsilateral phase variability, and upper intralimb phase variability), and the equivalence bound was set at Cohen’s d=0.36, estimated from the maximum bilateral difference in change in arm swing ROM due to DT in healthy older adults (Plate et al., 2015, tbl. 1).

Effect sizes were calculated using Cohen’s dav with a Hedge’s g correction, noted as gav (Cumming, 2011; Lakens, 2013). A sensitivity power analysis for a two-tailed t-test was conducted in G*Power (Faul et al., 2007) to find that the minimum detectable effect size with 80% power is *d*_*z*_ = 0.72 when *α* = .05 and the sample size is 17. A sensitivity power analysis for a TOST equivalence test using the TOSTER package in R showed that the minimum bounds that would be rejectable with 80% power is *d* = .90 when *α* = .05 and the sample size is 17. Arm swing for a given shoulder was treated as functionally absent when ROM was less than 5 deg. No meaningful coordination was expected between a shoulder with functionally absent arm swing and either the ipsilateral hip or the contralateral shoulder; therefore, the ipsilateral and intralimb CRP variables for affected subjects/shoulders were removed as outliers—3 subjects at most, depending on the variable. Circular statistics (circular mean and standard deviations) were used for the variables with angular units (Fisher, 1993). All data reduction and statistical analyses were performed with the Julia language using open-source libraries and code (Bezanson et al., 2017; Hill & Nantel, 2023).

## Results

Participant demographics are reported in Table 1. Twenty subjects who met the inclusion criteria were recruited. Two participants with severe dyskinesia were excluded from this study, and a third participant was excluded for moving their hands while talking during a significant portion of a trial; both behaviors produce movements (Jankovic, 2005) which are disruptive to the normal coordination patterns of steady state gait. Subjects were diagnosed with PD an average of 7.4 ± 4.5 years prior to study participation.

**Table 1.** Participant demographics.

| Characteristic | Mean ± SD | Range |
| --- | --- | --- |
| Sex (n) | 11 M, 6 F |  |
| Age (years) | $64.8 \pm 7.7$ | 48-79 |
| Height (cm) | $173.3 \pm 7.6$ | 165-188 |
| Weight (kg) | $74.2 \pm 19.9$ | 52-128 |
| Handedness | 14 R, 3 L |  |
| More affected side | 9 R, 8 L |  |
| UPDRS III | $10.2 \pm 5.3$ | 0-20 |
| Freezers (n) | 5 |  |
| Fallers (n, <1yr) | 10 |  |

Single and dual task results were calculated using an average of 139 ± 19 and 74 ± 6 steps, respectively (dual task trial length was shorter than the single task trial). Preferred gait speed was 1.0 ± 0.2 m/s among subjects.

### Unilateral effects of DT

Spatial coordination results are reported in Table 2. The LA shoulder ROM and peak flexion decreased by 1.5 deg 95% CI [−2.8,−0.2] and 1.1 deg [−2.1,−0.1], respectively, during dual task compared to single task walking. Temporal coordination results are reported in Table 3; no significant differences were detected (p>.061).

**Table 2.** Spatial coordination in the more and less affected sides during single and dual task.

| Variable | | Single task | Dual task | t-test | p-value | $g_{av}$ |
| --- | --- | --- | --- | --- | --- | --- |
| Shoulder ROM (deg) | LA | $21.2 \pm 13.3$ | $19.7 \pm 12.4$ | $t(16) = -2.44$ | **0.027** | -0.12 |
| | MA | $17.9 \pm 12.6$ | $19.7 \pm 14.8$ | $t(16) = 0.69$ | 0.501 | 0.13 |
| Shoulder ROM COV (%) | LA | $17.2 \pm 9.1$ | $25.0 \pm 38.5$ | $t(16) = 1.01$ | 0.329 | 0.32 |
| | MA | $22.1 \pm 14.8$ | $16.4 \pm 6.6$ | $t(16) = -1.55$ | 0.141 | -0.51 |
| Shoulder peak flexion (deg) | LA | $15.3 \pm 8.6$ | $14.2 \pm 8.2$ | $t(16) = -2.41$ | **0.028** | 0.47 |
| | MA | $13.3 \pm 7.7$ | $14.1 \pm 9.3$ | $t(16) = 0.48$ | 0.64 | 0.05 |
| Hip ROM (deg) | LA | 38.5 ± 4.1 | 37.6 ± 3.9 | t(16)=-1.61 | 0.127 | -0.22 |
|  | MA | 37.6 ± 5.1 | 37.6 ± 5.3 | t(16)=0.14 | 0.892 | 0.01 |
| Hip ROM COV (%) | LA | 4.2 ± 1.4 | 4.7 ± 1.8 | t(16)=1.82 | 0.087 | 0.27 |
|  | MA | 4.5 ± 1.6 | 4.6 ± 1.8 | t(16)=1.12 | 0.278 | 0.10 |
| Hip peak flexion (deg) | LA | 22.7 ± 3.9 | 22.4 ± 4.4 | t(16)=-0.92 | 0.374 | -0.01 |
|  | MA | 20.4 ± 5.1 | 20.5 ± 5.0 | t(16)=0.10 | 0.923 | -0.18 |

Note. ROM = range of motion, MA/LA = more/less affected side, $g_{av}$ = Cohen's $d_{av}$ effect size corrected with Hedge's $g$ .

**Table 3.**
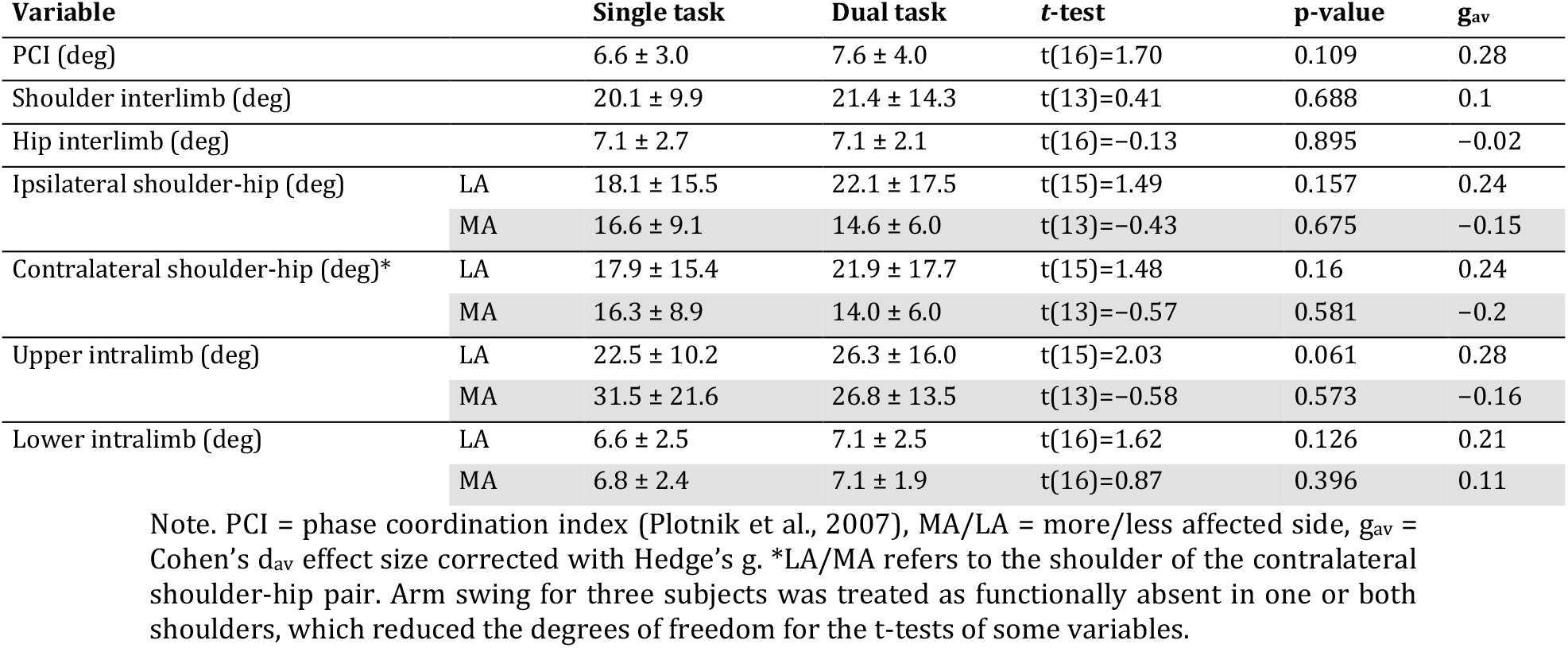
Temporal coordination (PCI and phase variability) in and between the more and less affected sides during single and dual task.

| Variable | | Single task | Dual task | t-test | p-value | $g_{av}$ |
| --- | --- | --- | --- | --- | --- | --- |
| PCI (deg) |  | 6.6 ± 3.0 | 7.6 ± 4.0 | t(16)=1.70 | 0.109 | 0.28 |
| Shoulder interlimb (deg) |  | 20.1 ± 9.9 | 21.4 ± 14.3 | t(13)=0.41 | 0.688 | 0.1 |
| Hip interlimb (deg) |  | 7.1 ± 2.7 | 7.1 ± 2.1 | t(16)=-0.13 | 0.895 | -0.02 |
| Ipsilateral shoulder-hip (deg) | LA | 18.1 ± 15.5 | 22.1 ± 17.5 | t(15)=1.49 | 0.157 | 0.24 |
|  | MA | 16.6 ± 9.1 | 14.6 ± 6.0 | t(13)=-0.43 | 0.675 | -0.15 |
| Contralateral shoulder-hip (deg)* | LA | 17.9 ± 15.4 | 21.9 ± 17.7 | t(15)=1.48 | 0.16 | 0.24 |
|  | MA | 16.3 ± 8.9 | 14.0 ± 6.0 | t(13)=-0.57 | 0.581 | -0.2 |
| Upper intralimb (deg) | LA | 22.5 ± 10.2 | 26.3 ± 16.0 | t(15)=2.03 | 0.061 | 0.28 |
|  | MA | 31.5 ± 21.6 | 26.8 ± 13.5 | t(13)=-0.58 | 0.573 | -0.16 |
| Lower intralimb (deg) | LA | 6.6 ± 2.5 | 7.1 ± 2.5 | t(16)=1.62 | 0.126 | 0.21 |
|  | MA | 6.8 ± 2.4 | 7.1 ± 1.9 | t(16)=0.87 | 0.396 | 0.11 |

Note. PCI = phase coordination index (Plotnik et al., 2007), MA/LA = more/less affected side, $g_{av}$ = Cohen's $d_{av}$ effect size corrected with Hedge's $g$ . \*LA/MA refers to the shoulder of the contralateral shoulder-hip pair. Arm swing for three subjects was treated as functionally absent in one or both shoulders, which reduced the degrees of freedom for the t-tests of some variables.

### Bilateral difference in DTC

Equivalence tests for difference in DTC between sides are reported in Table 4. Hip ROM DTC was significantly different between sides, *M* = −0.967°, and was not equivalent (*p* = .88), preventing a conclusion of equivalence to a healthy older adult population (See Fig. 1). Lower intralimb phase variability DTC was not significantly different between sides, and was statistically equivalent (*p* = .004).

**Table 4.** Tests of equivalence in DTC between sides.

| DTC Variable | LA side | MA side | 95% CI | Difference t-test | p-value | TOST p-value | $g_{av}$ |
| --- | --- | --- | --- | --- | --- | --- | --- |
| Shoulder ROM (deg) | -1.5 ± 2.6 | 1.8 ± 11 | [-8.8, 2.2] | t(16)=-1.26 | 0.225 | 0.467 | -0.49 |
| Shoulder ROM COV (%) | 7.8 ± 32 | -5.6 ± 15 | [-4.1, 31] | t(16)=1.62 | 0.125 | 0.808 | 0.56 |
| Shoulder peak flexion (deg) | -1.1 ± 2 | 0.78 ± 6.8 | [-5.5, 1.7] | t(16)=-1.13 | 0.276 | 0.177 | -0.42 |
| Hip ROM (deg) | -0.92 ± 2.3 | 0.051 ± 1.5 | [-1.8, -0.15] | t(16)=-2.52 | **0.023** | 0.88 | -0.49 |
| Hip ROM COV (%) | 0.45 ± 1 | 0.16 ± 0.6 | [-0.23, 0.81] | t(16)=1.17 | 0.26 | 0.378 | 0.35 |
| Hip peak flexion (deg) | -0.21 ± 0.96 | 0.027 ± 1.2 | [-0.96, 0.49] | t(16)=-0.69 | 0.502 | 0.223 | -0.22 |
| Ipsilateral shoulder-hip (deg) | 5 ± 11 | -1.1 ± 9.6 | [-1.9, 14] | t(13)=1.64 | 0.124 | 0.614 | 0.59 |
| Upper intralimb (deg) | 2.1 ± 5.2 | -2.8 ± 18 | [-6.4, 16] | t(13)=0.93 | 0.369 | 0.342 | 0.41 |
| Lower intralimb (deg) | 0.53 ± 1.3 | 0.24 ± 1.1 | [-0.11, 0.68] | t(16)=1.53 | 0.146 | **0.004** | 0.22 |

Note. MA/LA = more/less affected side, $g_{av}$ = Cohen's $d_{av}$ effect size corrected with Hedge's $g$ . Arm swing for three subjects was treated as functionally absent in one or both shoulders, which reduced the degrees of freedom for the t-tests of some variables.

**Figure 1.**
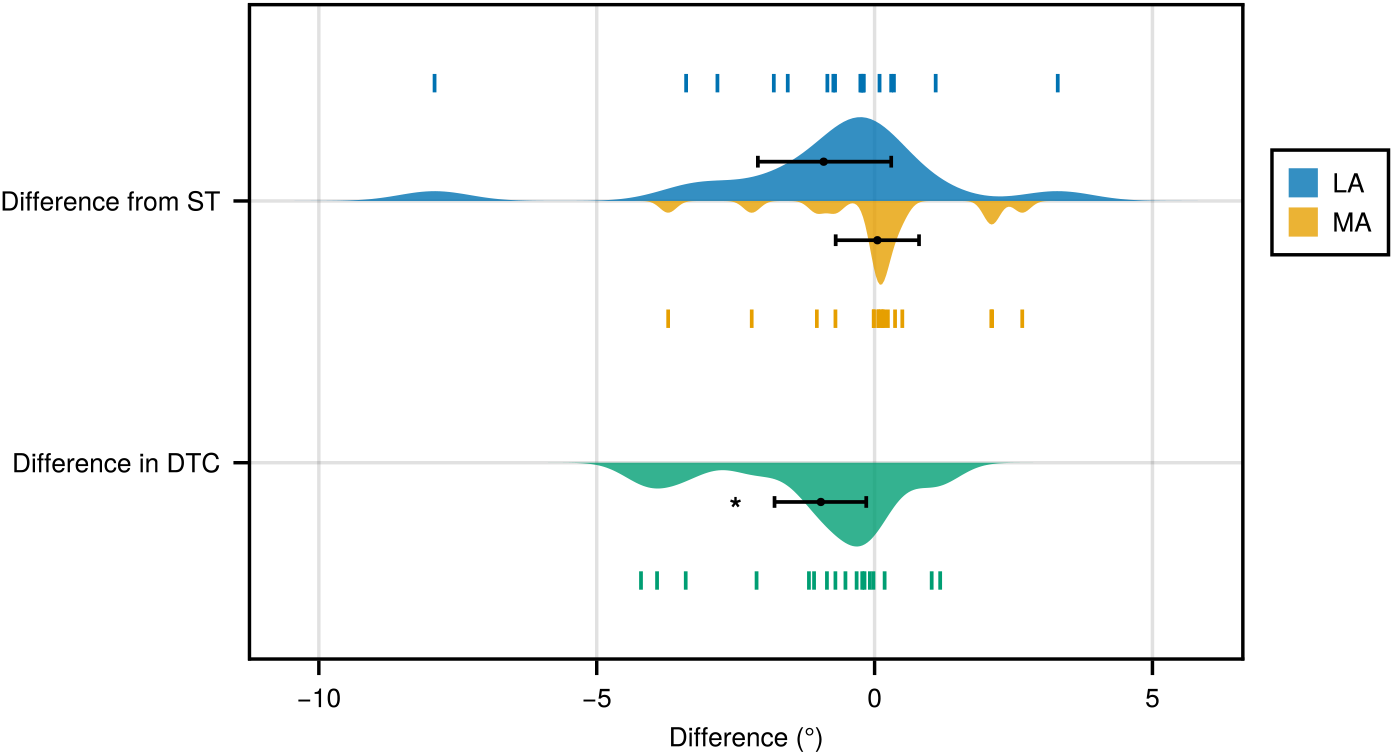
Distribution and CIs of changes in unilateral hip ROM due to dual-tasking. Error bars represent the 95% CIs. On top, the LA and MA hip ROM did not significantly change between single and dual task conditions. On bottom, the asterisk denotes that the difference in DTC for hip ROM was significantly different between LA and MA sides.

## Discussion

In this study, we found that few aspects of coordination are unilaterally affected by a visuospatial dual task in a cohort of mild to moderate PD. Furthermore, few aspects of coordination exhibit a significant bilaterally different sensitivity to dual tasking, however, most variables do not support a conclusion of equivalent sensitivity of coordination between the MA and LA sides. During the dual task, the LA shoulder ROM and peak flexion decreased compared to single task performance (Table 2), but the change in shoulder ROM and peak flexion due to dual tasking were not significantly different between the LA and MA sides. A significant difference was noted in hip ROM DTC between MA and LA sides, but neither side was affected by dual tasking. Lower intralimb phase variability was not affected by dual tasking and was not different and equivalent between sides at ±0.85° bounds.

Our results are generally inconclusive, as null findings for both t-tests and equivalence tests prevent meaningful conclusions about the presence or absence of meaningful (i.e. clinically relevant) effects. Reduced arm swing during dual tasking on the LA side is similar to previous studies (Mirelman et al., 2016; Baron et al., 2018). Additionally, gait asymmetry present in people with PD often results in LA ROM and coordination that are more similar to—but not always matching—healthy peers, while the MA side is less similar to healthy peers (Roggendorf et al., 2012; Roemmich et al., 2013). Qualitatively, the single task shoulder ROM in our data showed that the arm swing was fairly symmetric (Table 2), and slightly reduced on both sides compared to normative arm swing data for healthy older adults at similar gait speeds (Plate et al., 2015; Killeen et al., 2018).

However, we note that the absence of a significant effect of dual tasking on the MA shoulder ROM should not be interpreted as evidence for differing behaviors/responses to dual tasking between the MA and LA shoulder ROM, as Gelman and Stern (2006) have previously stated, “comparisons of the sort ‘X is statistically significant and Y is not’ can be misleading”. Indeed, we directly compared the unilateral DTC and found no difference between sides for shoulder ROM. Since equivalence at the tested bounds was not reached, there remains the possibility of a bilateral difference in DTC at an effect size that is smaller than we were powered to detect but too large to reject at our chosen equivalence bounds. Similarly, while there were no significant differences between single and dual task for unilateral hip ROM, there was a significant difference in DTC between sides; see Fig. Figure 1.

Lower intralimb phase variability was not affected by the dual task used in this study, and the difference in responses of each side to dual tasking was equivalent to estimated responses in healthy older adults. Different dual tasks have been found to have population specific effects (i.e. task interference) (Al-Yahya et al., 2011), and previous studies on gait and dual tasks in PD suggests that fallers and/or freezers may be more sensitive to dual task interference (Plotnik, Giladi, et al., 2011; Bekkers et al., 2018). More research is needed to determine whether such increased sensitivity for PD fallers or freezers would apply to the novel visuospatial task—intended to mimic an ecologically realistic scenario—that was used in this study.

### Limitations

Due to the absence of a control group, we are only able to quantify the response of a mild PD group to the DT, and are unable to discuss how responses to the DT in a group of healthy controls might differ from this group of people with PD. In addition, the participants in this study had mild to moderate PD, and the arm swing ROM and ROM asymmetry within our cohort is markedly different compared to previously reported PD cohorts (Roggendorf et al., 2012; Isaias et al., 2012; Sterling et al., 2015; Mirelman et al., 2016); it is unclear how coordination would respond in people with similarly mild PD—but with larger levels of arm swing asymmetry, or in people with more severe PD—which have more symmetric arm swing but reduced ROM (Roggendorf et al., 2012). Similarly, the severity of PD—in terms of UPDRS III scores—may be correlated with dual task interference, however the sparse range of UPDRS III values in our cohort prevented the use of statistical tests that might detect any such interactions.

Additionally, different tasks are known to have unique and specific effects on different aspects of gait which may limit the generalizability of our results (Al-Yahya et al., 2011; Rochester et al., 2014), and the visuospatial dual task in this study was simple (12/17 participants demonstrated perfect performance, and the remaining 5 participants responded to 10 ± 1.5 words out of 12). However, Baron et al. (2018) found that arm swing kinematics were sensitive to multiple common dual tasks.

Finally, treadmill walking is different from overground walking and may prevent some common responses in PD, such as reducing gait speed when distracted. However, our results are still informative about the nature of coordination deficits in PD and the methods they use to compensate for coordination deficits when optimally medicated and walking in this specific environment.

## Conclusion

Our results show that in a group of mild PD, a visuospatial dual task during gait contributes to decreased arm ROM on the LA side, but only hip ROM is differently affected by a dual task on the MA and LA sides. Furthermore, with the exception of coordination within the lower limbs, tests of equivalent sensitivity to dual tasking on the MA and LA sides were inconclusive. Despite the inconclusive tests, these results support cumulative science by providing reference data useful in calculating effect sizes for power analyses of future studies on the topic (Lakens, 2013). More research is needed to support or dispute the existence of differences in coordination between the MA and LA sides, and whether any differences are moderated by medication.

## Declarations

## Acknowledgements

Preprint version 3 of this article has been peer-reviewed and recommended by Peer Community In Health and Movement Sciences (https://doi.org/10.24072/pci.healthmovsci.100043; Ravi, 2024).

## Funding

This work was supported by the Natural Sciences and Engineering Research Council of Canada (https://www.nserc-crsng.gc.ca) [RGPIN-2016-04928 to J.N., RGPAS 493045-2016 to J.N.], and by the Ontario Ministry of Research, Innovation and Science (https://www.ontario.ca/page/early-researcher-awards) Early Researcher Award [ER 16-12-206 to J.N.]. The funders had no role in study design, data collection and analysis, decision to publish, or preparation of the manuscript.

## Competing interests

The authors have no relevant financial or non-financial interests to disclose.

## Ethics approval

This study was performed in accordance with the Declaration of Helsinki. Approval was granted by the Ottawa Health Science Network Research Ethics Board (No. 20170291-01H) and the University of Ottawa Office of Research Ethics and Integrity (No. A06-17-03).

## Consent to participate

Written informed consent was obtained from all study participants.

## Data and code availability

The software and data produced and analyzed for this study are openly available from the Zenodo data repository at https://doi.org/10.5281/zenodo.8364708 (Hill & Nantel, 2023)

## Author contributions

**Allen Hill:** Conceptualization, Methodology, Software, Validation, Formal Analysis, Investigation, Resources, Data Curation, Writing – Original Draft. **Julie Nantel:** Investigation, Resources, Writing – Review & Editing, Supervision, Project administration, Funding acquisition

## Notes

https://doi.org/10.5281/zenodo.6835766

### Competing Interest Statement

The authors have declared no competing interest.

### Summary of Updates

Manuscript has been updated to reflect its status as "Recommended" by the Peer Community in Health and Movement Sciences. A "Recommended" badge has been added to the cover page, as well as the addition of an acknowledgement of the recommendation process.

